# Imputing missing minimum inhibitory concentration (MIC) values for *Pseudomonas aeruginosa* strains with a Denoising AutoEncoder

**DOI:** 10.1101/2025.03.26.645597

**Authors:** Elsa Denakpo, Nicolas Dias, Johann D D Pitout, Thierry Naas, Dylan R Pillai, Flora Jay, Bogdan I. Iorga

## Abstract

*Pseudomonas aeruginosa* is a problematic pathogen with complex antibiotic resistance patterns. In clinical practice, minimum inhibitory concentration (MIC) tests typically focus on a limited subset of antibiotics, hindering a comprehensive assessment of a strain’s resistance profile. Here, we introduce MICFiller, a Denoising AutoEncoder (DAE) model designed to impute missing MIC values for 14 antibiotics in *Pseudomonas aeruginosa* within a specific dilution range by leveraging known MIC measurements for other antibiotics in the same strain. We evaluated the performance of DAE against two other commonly used methods: Multiple Imputation by Chained Equations (MICE) and simple median imputation. The DAE achieved the highest balanced 1-tier accuracy for most antibiotics, with performance closely matching that of MICE. MICFiller is freely accessible through a user-friendly web interface at http://iorgalab.org:4567/micfiller, offering clinicians a more complete view of a strain’s antibiotic resistance profile.

## Introduction

*Pseudomonas aeruginosa* is a clinically significant pathogen that poses a major challenge due to its complex and rapidly evolving antibiotic resistance mechanisms. In hospital settings, this organism is a frequent cause of serious infections, including pneumonia, bloodstream infections, and urinary tract infections. Effective treatment hinges on accurate antibiotic susceptibility testing, often assessed through minimum inhibitory concentration (MIC) measurements. The MIC is obtained by exposing bacteria to a range of antibiotic concentrations using serial dilutions until no visible growth is observed, thus identifying the lowest concentration capable of inhibiting bacterial proliferation.

Despite its clinical importance, MIC determination can be both labor-intensive and time-consuming. Consequently, many healthcare facilities routinely test a narrowed panel of antibiotics, rather than a more extensive array, to streamline resource use and speed up patient care decisions. However, focusing on a limited set of antibiotics can obscure a comprehensive understanding of a strain’s resistance profile. This is particularly problematic for *P. aeruginosa*, which can acquire or upregulate resistance mechanisms at an alarming rate. As a result, treating physicians may be left with incomplete susceptibility information, potentially leading to suboptimal therapeutic choices.

One promising approach to address this limitation is the application of data imputation algorithms. By leveraging available MIC measurements for a subset of antibiotics, these methods can estimate missing MIC values for other antibiotics, thereby filling critical gaps in the resistance profile. Not only could this yield a more complete picture of a strain’s susceptibility, but it might also reduce the need for additional time-consuming laboratory tests. Recently, advanced machine learning techniques—such as Denoising AutoEncoders—have shown particular promise for handling partially observed, high-dimensional datasets.

In this study, we systematically explore multiple imputation techniques for recovering missing MIC values in *P. aeruginosa*, encompassing both established methods such as Multiple Imputation by Chained Equations (MICE) and median-based strategies, as well as a novel Denoising AutoEncoder (DAE) approach. We begin by describing the dataset’s composition and clinical context, including the specific range of MIC measurements examined. We then detail the theoretical foundations and practical implementation of each method, highlighting key performance metrics and potential applications. Finally, we present and interpret our findings in the context of clinical decision-making, emphasizing how these imputation tools may be effectively integrated into laboratory workflows to enhance antibiotic susceptibility assessments.

## Dataset

This study is based on antimicrobial susceptibility data published by Stanton *et al*. [5] in 2022, which focused on 1,019 *Pseudomonas aeruginosa* isolates collected through the CDC’s (Centers for Disease Control and Prevention) Emerging Infections Program in the United States. All isolates underwent evaluation under broth microdilution conditions against 15 antibiotics commonly used to treat *P. aeruginosa* infections. The minimum inhibitory concentration (MIC) of each antibiotic was determined by exposing the bacterial isolate to a series of twofold dilutions, from which the lowest concentration inhibiting visible growth was recorded. This process yielded a distribution of MIC values spanning a clinically relevant range for each antibiotic. These antibiotics are categorized according to their families and mechanisms of action:

- *β*-lactams: aztreonam, cefepime, ceftazidime, ceftazidime-avibactam, ceftolozane-tazobactam, piperacillin-tazobactam, doripenem, meropenem, and imipenem
- Lipopeptides: colistin
- Fluoroquinolones: ciprofloxacin and levofloxacin
- Aminoglycosides: gentamicin, amikacin, and tobramycin

A principal goal of our investigation was to examine how MICs correlate with different antibiotics, with the understanding that correlations often indicate shared resistance mechanisms or overlapping targets. Figure 1 shows a Kendall *τ*_*c*_ rank correlation matrix of the log_2_ transformed MIC values for the 15 antibiotics, illustrating pairwise relationships for all 1,019 *P. aeruginosa* isolates. The transformation log_2_ helps normalize the discrete nature of MIC data, while Kendall *τ*_*c*_ is well suited for rank-based correlation, mitigating issues arising from ties in the observed values.

**Figure 1.**
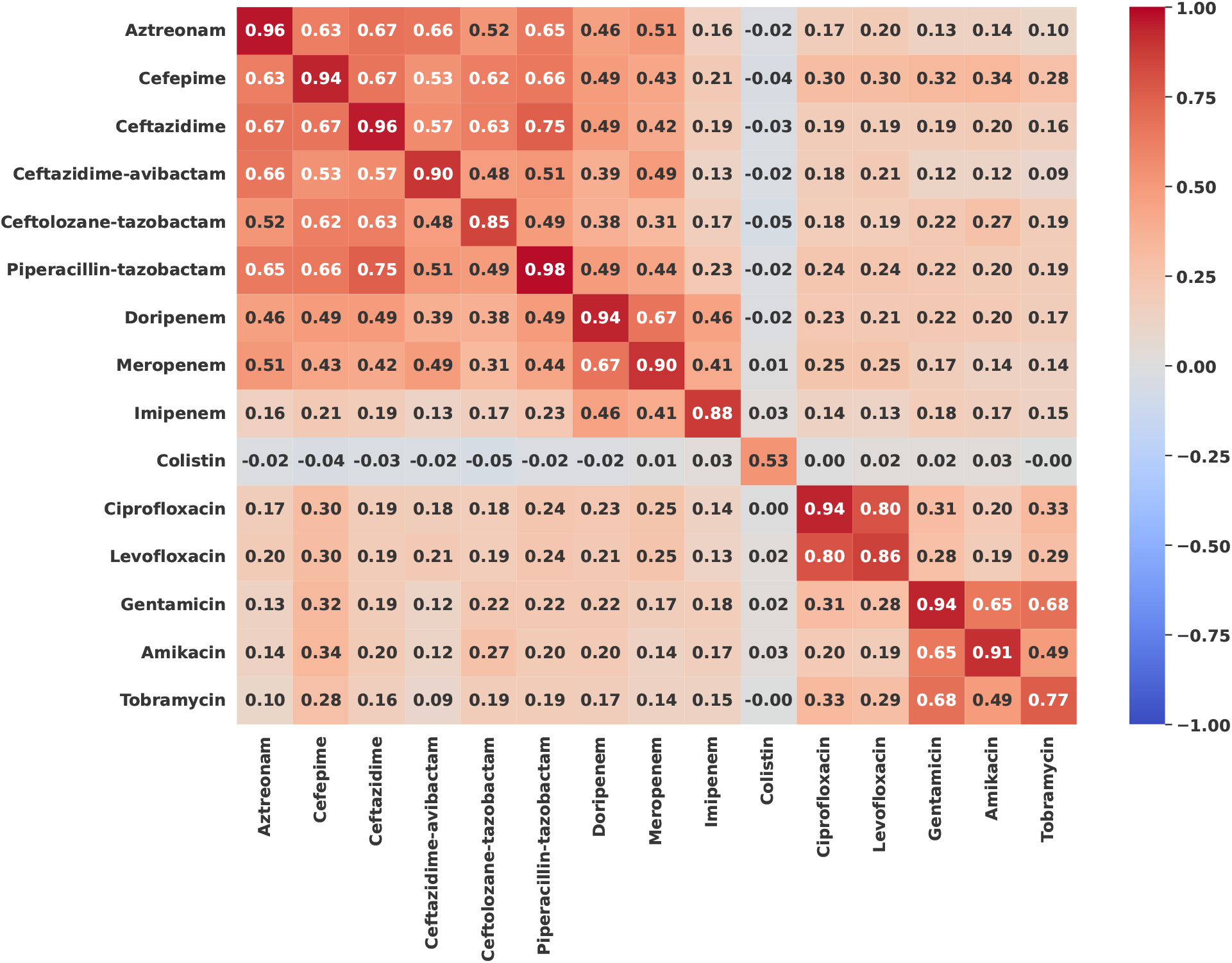
Kendall *τ*_*c*_ rank correlation correlation matrix between antibiotics log_2_-transformed MIC for 1019 *P. aeruginosa* strains from CDC’s dataset.

**Figure 2.**
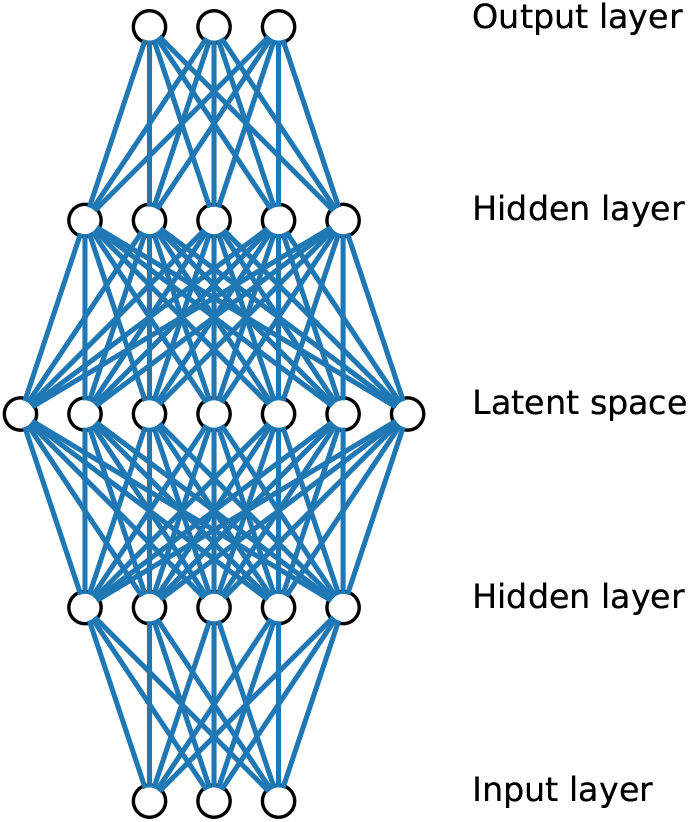
Overcomplete denoising autoencoder.

As shown in Figure 1, there are strong positive correlations among antibiotics within the same family. For example, most *β*-lactams cluster together, reflecting similar modes of action and comparable susceptibility profiles. However, imipenem stands out by correlating mainly with other carbapenems (doripenem and meropenem), suggesting that its specific mechanism differs from other *β*-lactams, leading to partially distinct resistance patterns. Beyond these intra-family correlations, cross-family relationships are also evident. For example, cefepime (a fourth-generation cephalosporin) shows moderate correlations with fluoroquinolones (ciprofloxacin and levofloxacin) and aminoglycosides (gentamicin, amikacin, and tobramycin). Similarly, tobramycin exhibits correlations with fluoroquinolones, hinting at potential partially overlapping resistance mechanisms between these drug classes.

In contrast, colistin does not show significant correlations with other antibiotics, possibly because its mode of action, membrane disruption, differs substantially from the targets of *β*-lactams, fluoroquinolones and aminoglycosides. In addition, colistin is often reserved for highly drug-resistant infections, so its MIC values may reflect a narrower subset of clinical scenarios. Given these differences, we exclude colistin from further analyses to focus on the remaining 14 antibiotics that exhibit more interconnected resistance profiles.

Together, these observed correlations among MIC values underscore the promise of leveraging known susceptibility information for certain antibiotics to infer missing MIC values for others. In the forthcoming sections, we detail a range of imputation methods and evaluate how accurately they can fill these gaps, ultimately aiming to provide a more comprehensive antibiotic resistance profile for each *P. aeruginosa* strain.

## Imputation of Missing Values

Imputation methods are designed to replace missing values in a dataset with reasonable estimates, thereby allowing downstream analyses to be performed without discarding incomplete records. These techniques span a broad range of complexity, from simple univariate strategies to more advanced regression-based or machine learning approaches. In this work, we focus on three methods that exemplify different levels of complexity and modeling assumptions. Each of these three approaches offers advantages and limitations in terms of computational complexity, interpretability, and the degree to which they incorporate relationships among variables.

### Univariate Imputation Method

Univariate imputation considers only the variable with missing entries when assigning replacement values. A straightforward implementation involves substituting each missing value with a constant, such as a mean or median computed from the observed data. In some cases, one may assume a specific parametric form (e.g., a normal distribution) and draw imputations according to estimated parameters (mean and variance). The main advantage of univariate approaches is their ease of implementation and low computational overhead. However, they do not capture multivariate dependencies among different variables, which can result in biased estimates if missingness is systematically related to other attributes.

### Multiple Imputation by Chained Equations (MICE)

Multiple Imputation by Chained Equations (MICE) [1] is a more sophisticated, multivariate approach. Initially, each variable with missing values is given a naive fill (e.g., a simple mean or median imputation). Subsequently, the variables are iteratively updated according to a predefined order: each variable with missing data is modeled as a function of the other variables, using only the observed and currently imputed values as inputs. Once a pass through all variables has been completed, a single “cycle” is considered finished.

Multiple cycles can then be performed, allowing the imputed values from previous steps to improve the model for the next iteration. This iterative procedure helps the algorithm converge toward imputations that better reflect the underlying correlations among variables. Although MICE can yield robust estimates of missing data, its computational cost grows with the number of cycles and the complexity of the chosen models. Careful consideration of convergence criteria and the choice of predictors for each variable is therefore crucial.

### Denoising AutoEncoder

An AutoEncoder is a neural network trained to slearn an identity mapping from input to output through a lower-dimensional (or sometimes higher-dimensional) latent space. It comprises two main components: an encoder, which projects the input into the latent space, and a decoder, which reconstructs the original input from this latent representation. A Denoising AutoEncoder (DAE) [6] extends this concept by learning to correct noise-corrupted inputs.

In the context of missing data, the absent entries can be treated as “noise.” During training, any missing values in the input variables must be filled with a placeholder (e.g., a constant) so that the network can process the entire feature set. The DAE then attempts to reconstruct the “true” values at the output layer. This approach is particularly useful when relationships among variables are nonlinear or when interactions are complex, as neural networks can automatically learn high-level abstractions.

Several recent studies indicate that using an overcomplete denoising autoencoder (DAE) can be more effective for noise reduction than an undercomplete autoencoder. In an overcomplete DAE the latent space has a higher dimension than the input space. This increased capacity allows the network to capture fine-grained features and subtle details that are critical for reconstructing a clean signal. When combined with regularization techniques (e.g., sparsity constraints), the overcomplete architecture learns to filter out noise without simply memorizing the identity mapping. For example, in the context of speech recognition, Kabil *et al*. [4] compared shallow undercomplete autoencoders, whose behavior is similar to principal component analysis (PCA) under certain conditions, with deeper, sparser overcomplete architectures. Their results demonstrated that sparse overcomplete autoencoders provided a clear advantage in reconstructing noisy Mel-Frequency Cepstral Coefficient (MFCC) features, thereby yielding improvements in recognition accuracy. Similarly, in skeleton-based action recognition, Guo *et al*. [3] proposed an overcomplete graph convolutional denoising autoencoder (GCDAE) that maintained more channels in its latent representation. This design enabled the model to preserve subtle spatial details (e.g., precise joint positions) that are often lost when the input is compressed into a smaller latent space in undercomplete models. Consequently, the overcomplete GCDAE achieved enhanced robustness against various types of noise and occlusion.

## Methodology

Our overall methodology includes the simulation of missing values in the dataset, the encoding strategy for MIC data, and the training procedures for each imputation approach. We also introduce the performance metric used to compare model outputs.

### Missingness Simulation

To investigate the ability of different imputation methods to predict missing MIC values, we artificially introduce missingness into the original dataset under the assumption of Missing At Random (MAR). Rather than maintaining a fixed rate of missingness, we vary the proportion of missing entries across different strains to create a more versatile training environment. Specifically, we use the following procedure:

- For each row (i.e., each *P. aeruginosa* strain), we randomly select one or more MICs to mark as missing, subject to two constraints: (1) at least one MIC in every antibiotic family must remain unmasked, and (2) at least one MIC in each row must be missing.
- We repeat this procedure *n* times over the entire dataset, creating multiple augmented copies of each strain with different missingness patterns.
- Duplicate rows generated through this augmentation are removed, and the resulting dataset is shuffled to prevent any ordering biases.

By applying various missingness rates, we train models that can adapt to a wide range of clinical scenarios, from relatively sparse information (many missing MICs) to more detailed phenotypic data (fewer missing MICs).

### MICs Encoding

Minimum Inhibitory Concentrations (MICs) are typically measured via two-fold dilutions in a bounded range. As a result, MIC values may be reported using three possible notations:

- <= *x*, indicating that the MIC is below or equal to the lower bound of the tested range,
- == *x*, indicating that the MIC falls within the tested range and is measured exactly as *x*,
- > *x*, indicating that the MIC exceeds the upper bound of the tested range.

Since machine learning algorithms require numeric inputs, we adopt the following encoding scheme:

1. If the MIC is reported as <= *x*, we replace it with the lower bound value *x*.
2. If the MIC is reported as == *x*, we keep it as *x*.
3. If the MIC is reported as > *x*, we replace it with the next value in the two-fold dilution series just above *x*.

After applying these rules, each MIC is transformed into a single numeric value. We then apply a log_2_ transformation, to have the same interval between each two consecutive MIC values. Finally, we perform min-max scaling to normalize the MIC data for each antibiotic into the [0, 1] range. This step provides a consistent input domain for training, reflecting the fact that each antibiotic has its own intrinsic dilution bounds.

### Training Process

Our core dataset consists of 1,019 *P. aeruginosa* strains and 14 antibiotics (features), with each antibiotic MIC acting both as a feature (predictor) and as a target (to be imputed). Below, we outline the training protocol for each of the imputation methods considered.

#### Data Splitting and Simulation

We apply the missingness simulation separately on the original dataset, repeating our missingness procedure *n* = 1,000 times. We then randomly partition the resulting dataset into a training set (80%) and a test set (20%). This yields 654,480 augmented examples in the training set and 163,972 examples in the test set. Models are then trained on the augmented training set to increase their robustness to different patterns and rates of missingness.

When methods like MICE or DAE require hyperparameter tuning through cross-validation, we simulate missingness in each fold independently, preserving the original stratification. For stratification, we choose the antibiotic with the most imbalanced MIC distribution as the stratification variable, ensuring that each fold maintains a representative distribution of challenging samples.

#### Univariate Imputation

For the univariate imputer, we replace each missing entry with the median MIC of the corresponding antibiotic. This median is computed from the training set *before* any missingness or augmentation is introduced, ensuring that we use a single fixed value across all augmented training and test examples. This simple approach serves as a baseline for more advanced imputation techniques.

#### MICE Imputation

We employ Multiple Imputation by Chained Equations (MICE) [1] with Random Forest estimators to model each antibiotic MIC as a function of the other antibiotics. The missing values are initially filled with their respective medians. During training, MICE iterates over all variables in a predefined order, taken from left to right in the correlation matrix (see Fig. 1), and fits a Random Forest using the observed or imputed values from the other variables. A grid search with 5-fold cross-validation is used to tune the maximum number of cycles (1, 2, 3, 4 or 5) and the Random Forest min *samples split* parameter (0.003125, 0.00625, 0.0125, 0.025 and 0.05). The best model identified by cross-validation is then retrained on the entire training set and evaluated on the test set. In our experiments, a total of 100 trees is used in each Random Forest, and the maximum number of features for each split is set to the square root of the total features.

#### Denoising AutoEncoder

For the Denoising AutoEncoder (DAE), we treat missing MIC values as “noise” by initially replacing them with −1. The network is designed with an input layer of 14 neurons (one per antibiotic), followed by a hidden layer with *H*_1_ neurons, a dropout layer, a hidden layer (the latent space) with *H*_2_ neurons, another hidden layer with *H*_3_ neurons, and an output layer of 14 neurons with a sigmoid activation.

We adopt an “overcomplete” strategy, where the dimensionality in the hidden layers may exceed that of the input, which is known to encourage better reconstruction when the data contains missing entries [2]. The number of units in each hidden layer follows a pattern determined by *θ*, which we vary in the set {5, 10, 15, 20, 25, 30, 35, 40, 45, 50, 55, 60}. Similarly, the dropout rate is tuned among {0.0, 0.1, 0.2, 0.3} using a 5-fold cross-validation. Rectified Linear Unit (ReLU) activation is applied to hidden layers, while the output layer uses a sigmoid function with Mean Squared Error as the reconstruction loss. We train this network using the Adam optimizer with exponential decay at a rate of *e*^*−*0.2^, starting from a learning rate of 10^*−*2^. Each model is trained for 20 epochs with a batch size of 128.

### Performance Metric: Balanced 1-Tier Accuracy

We evaluate each model’s imputation performance using the mean of the balanced 1-tier accuracy across all antibiotics. Let *A* be the set of antibiotics and *C*_*a*_ be the set of possible MIC classes for antibiotic *a* ∈ *A*. We first compute the 1-tier accuracy for each class *c* ∈ *C*_*a*_, defined as the proportion of predicted values that fall within ± 1 two-fold dilution factor of the true class *c*. The overall balanced 1-tier accuracy for each antibiotic is then the average of its class-specific accuracies. Finally, we take the mean across all antibiotics:

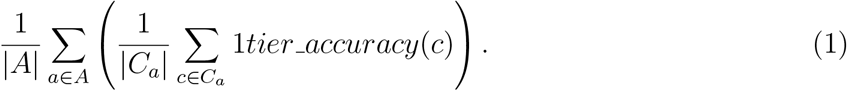

A higher balanced 1-tier accuracy indicates better agreement between imputed and true MIC values, giving equal weight to classes that are underrepresented in the dataset. This metric is especially pertinent in clinical contexts where rare resistance phenotypes must be accurately identified.

Overall, this methodology, combining missingness simulation, MIC encoding, data splitting, cross-validation for model selection, and a clinically relevant performance metric, allows us to comprehensively assess how well different imputation techniques can predict missing MIC values in *P. aeruginosa*.

## Results

The DAE gives better results compared with the MICE for some antibiotics, but they are very close for most antibiotics. To further analyze the DAE results, we plotted the performances on the missing values for each class within each antibiotic, when only one value is missing by row or in the extreme case when 11 values are missing by row (Fig 5, 6, 3, 4).

**Figure 3.**
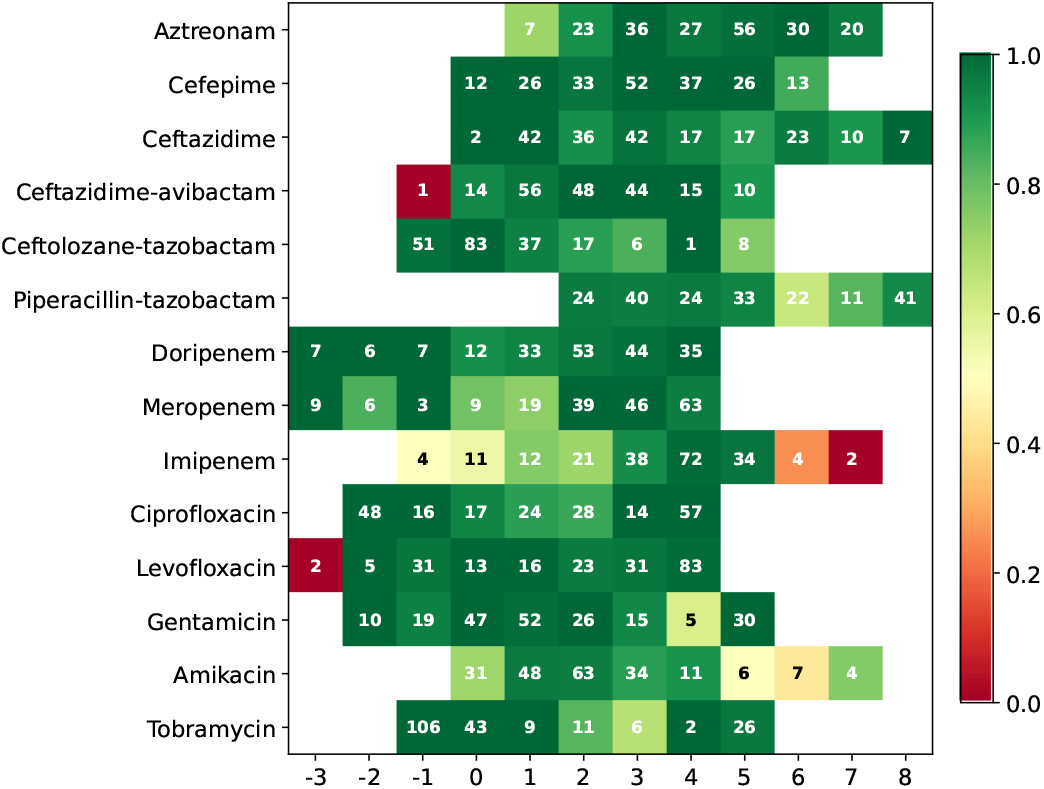
Recall on rows with 1 missing value: Each case represent an antibiotic and a MIC value. The number *n* in each case represent the number of strains which satisfy the following condition: of all the rows with only 1 missing values, how many with this missing antibiotic should belong to this specific class. The color degree within each case represents the DAE 1-tier accuracy values computed for those *n* strains

**Figure 4.**
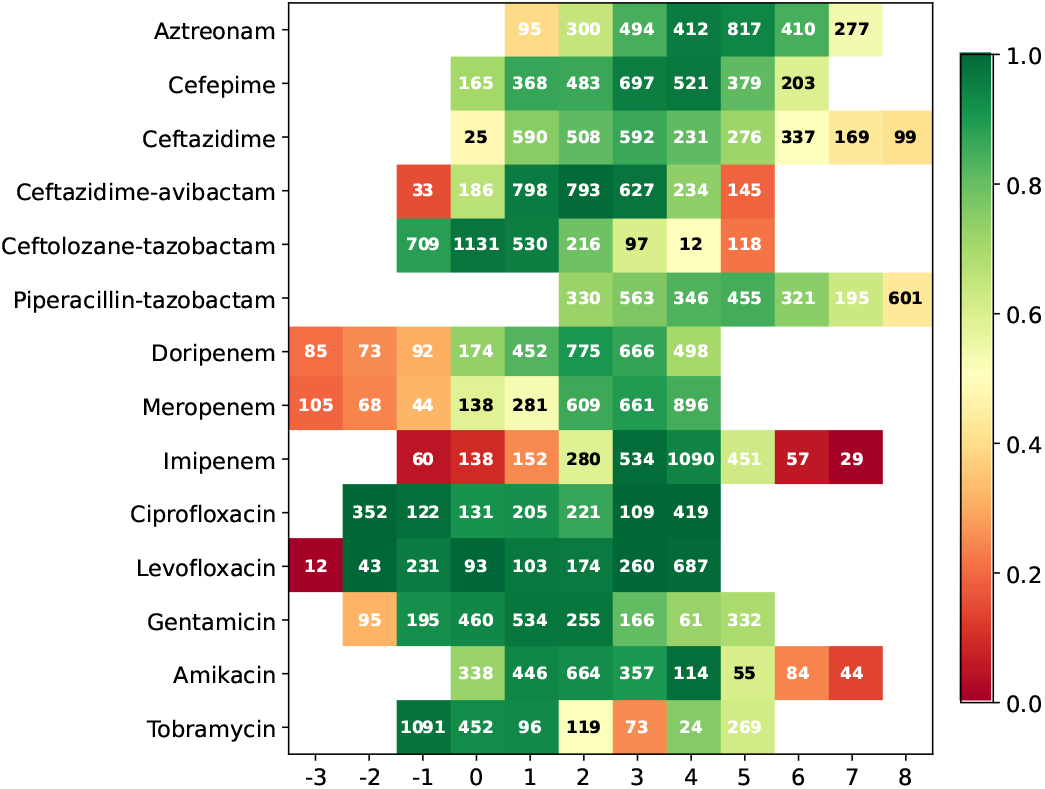
Recall on rows with 11 missing values: Each case represent an antibiotic and a MIC value. The number *n* in each case represent the number of strains which satisfy the following condition: of all the rows with only 11 missing values, how many with this missing antibiotic should belong to this specific class. The color degree within each case represents the DAE 1-tier accuracy values computed for those *n* strains

**Figure 5.**
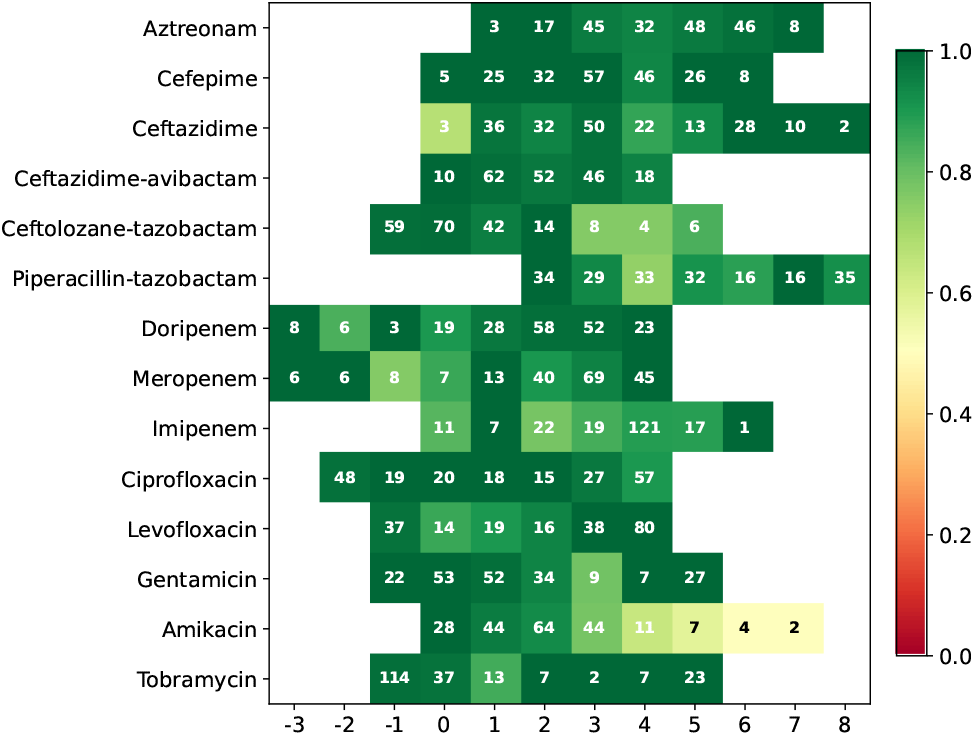
Precision on rows with 1 missing value: Each case represent an antibiotic and a MIC value. The number *n* in each case represent the number of strains which satisfy the following condition: of all the rows with only 1 missing values, how many have this missing antibiotic and have been predicted belonging to this specific class or the class before and after. The color degree within each case represents the precision values computed for those *n* strains

**Figure 6.**
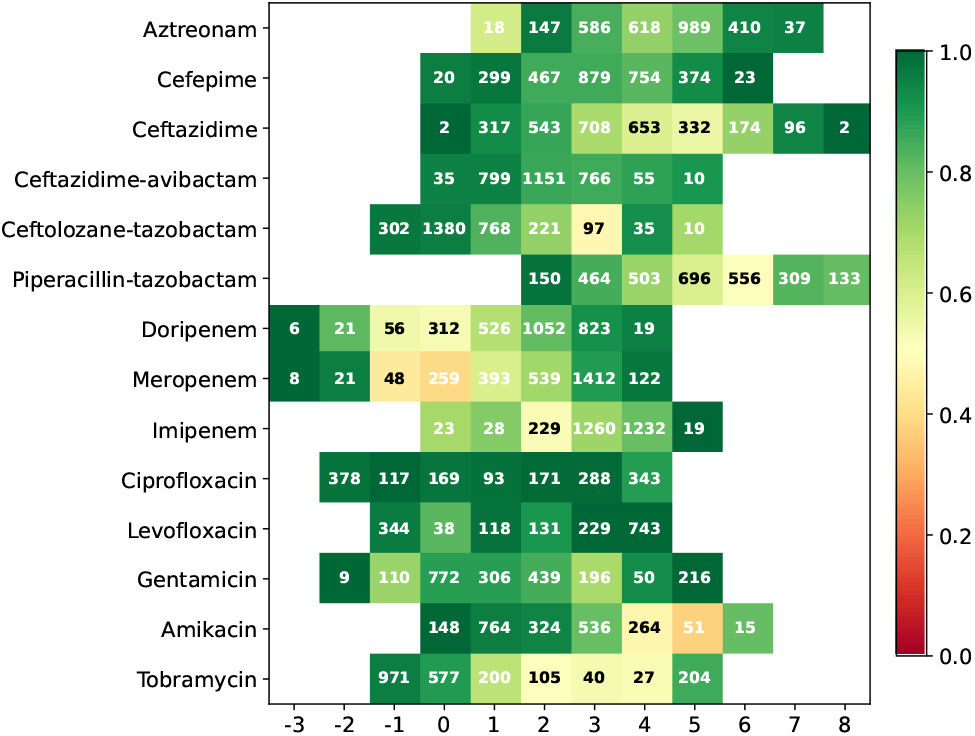
Precision on rows with 11 missing values: Each case represent an antibiotic and a MIC value. The number *n* in each case represent the number of strains which satisfy the following condition: of all the rows with only 11 missing values, how many have this missing antibiotic and have been predicted belonging to this specific class or the class before and after. The color degree within each case represents the precision values computed for those *n* strains

The algorithm works well when the number of missing values is lower. However, for some specific MICs, even with only one missing value, the accuracy is very low. That can be explained by the unequal distribution of MIC within each antibiotic which can be challenging for predicting the minority classes.

**Table 1.** Prediction results on the test set for the median imputer, MICE and DAE showing the mean absolute error and the balanced 1-tier accuracy on missing values.

|  | SI MAE | MICE MAE | DAE MAE | SI b1-tier | MICE b1-tier | DAE b1-tier |
| --- | --- | --- | --- | --- | --- | --- |
| Aztreonam | 1.384 | 0.68 | **0.664** | 0.429 | 0.849 | **0.859** |
| Cefepime | 1.255 | 0.561 | **0.559** | 0.429 | 0.912 | **0.92** |
| Ceftazidime | 1.634 | **0.752** | 0.765 | 0.333 | **0.828** | 0.826 |
| Ceftazidime-avibactam | 0.983 | 0.565 | **0.55** | 0.429 | 0.789 | **0.799** |
| Ceftolozane-tazobactam | 0.896 | **0.523** | 0.558 | 0.429 | **0.845** | 0.84 |
| Piperacillin-tazobactam | 1.77 | **0.836** | 0.841 | 0.429 | 0.794 | **0.8** |
| Doripenem | 1.265 | **0.76** | 0.774 | 0.375 | 0.757 | **0.773** |
| Meropenem | 1.346 | **0.749** | 0.786 | 0.375 | 0.695 | **0.715** |
| Imipenem | 1.119 | **0.924** | 0.933 | 0.333 | **0.529** | 0.525 |
| Ciprofloxacin | 2.048 | 0.515 | **0.39** | 0.429 | **0.956** | 0.95 |
| Levofloxacin | 1.659 | 0.387 | **0.306** | 0.375 | 0.854 | **0.86** |
| Gentamicin | 1.5 | 0.651 | **0.617** | 0.375 | 0.838 | **0.841** |
| Amikacin | 1.132 | **0.711** | 0.732 | 0.375 | 0.734 | **0.738** |
| Tobramycin | 1.395 | 0.537 | **0.504** | 0.286 | 0.774 | **0.777** |

## Conclusion

In this study, we investigated three different approaches to impute missing minimum inhibitory concentration (MIC) values in *Pseudomonas aeruginosa*: a univariate median imputation method, Multiple Imputation by Chained Equations (MICE), and a Denoising AutoEncoder (DAE). Through a controlled missingness simulation and comprehensive evaluation on a large dataset of clinically relevant *P. aeruginosa* isolates, we observed that both MICE and DAE substantially outperformed the univariate strategy across most antibiotics. Although the DAE demonstrated marginal improvements over MICE for several compounds, their overall performance was comparable, underscoring the robustness of both multivariate approaches.

A key finding was that imputation accuracy declined with increasing rates of missingness, which indicates the importance of incorporating as much phenotypic information as possible. Furthermore, we noted that the uneven distribution of MIC values, a common characteristic of clinical datasets, posed additional challenges for minority classes. Nevertheless, the strong correlations among antibiotics within and across families offered an effective basis for imputing missing MICs.

Our results highlight the value of using data-driven approaches for imputing incomplete susceptibility profiles in *P. aeruginosa*. Such methods have practical relevance in both research and clinical laboratories, where resources or time constraints may limit routine testing of all antibiotics. Future work could explore the application of this methodology to other bacterial species, ensemble methods that combine the strengths of multiple imputation algorithms, as well as investigate alternative neural network architectures tailored to the hierarchical nature of antimicrobial susceptibility data. Ultimately, improving MIC prediction through robust imputation techniques can offer a more complete picture of bacterial resistance, facilitating better-informed therapeutic decisions and contributing to more effective antibiotic stewardship.

## Acknowledgments

This work was supported, in part, by grants from the PPR Antibioresistance (ANR-20-PAMR-0010) and UDOPIA (ANR-20-THIA-0013).

## Notes

### Competing Interest Statement

The authors have declared no competing interest.

