## Supplementary information file for "Imputing missing minimum inhibitory concentration (MIC) values for *Pseudomonas aeruginosa* strains with a Denoising AutoEncoder"

### MIC distribution of antibiotics

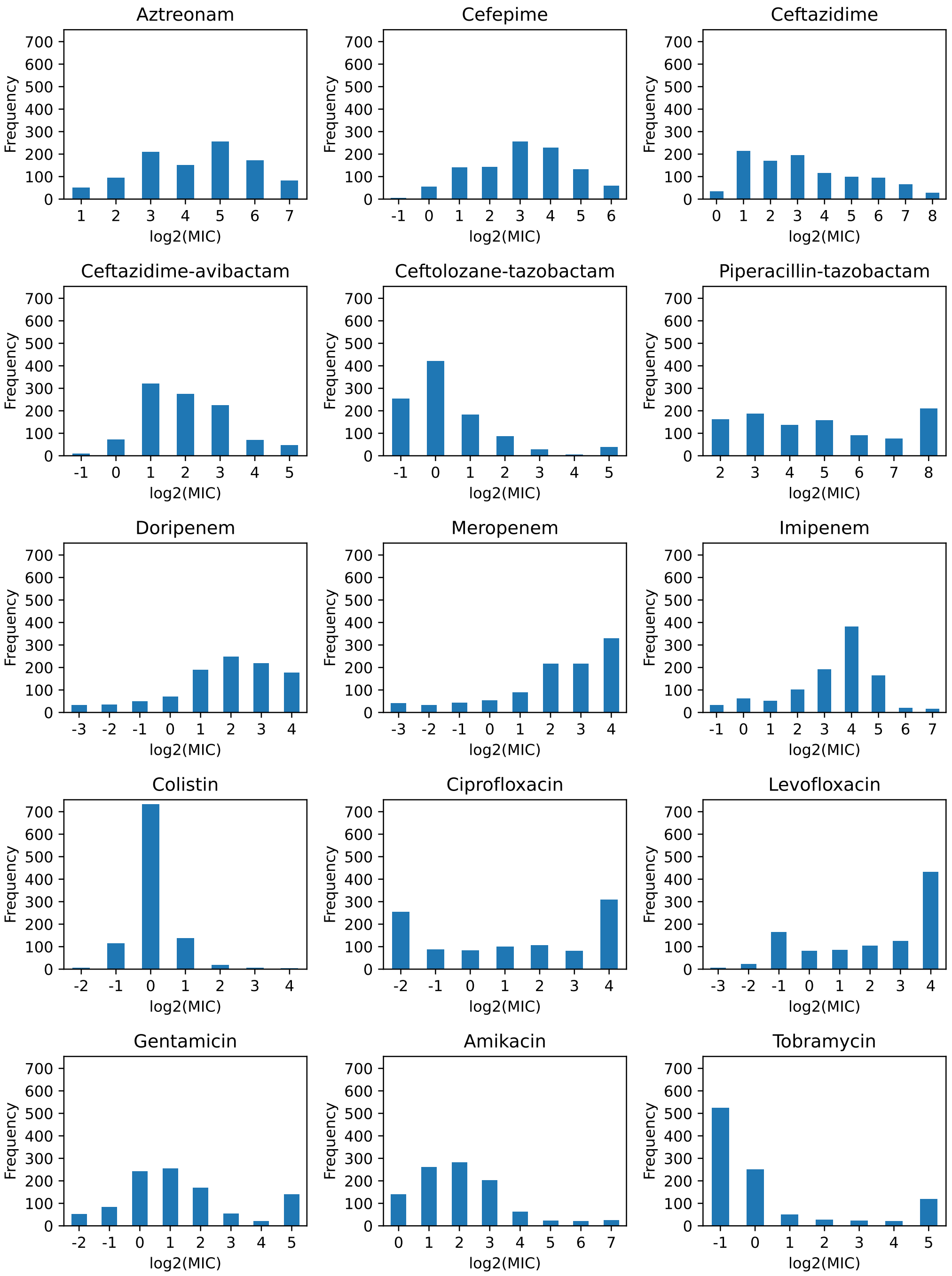


**Figure S1.** MIC distribution of 15 antibiotics from the dataset used in this study

### MICE performances

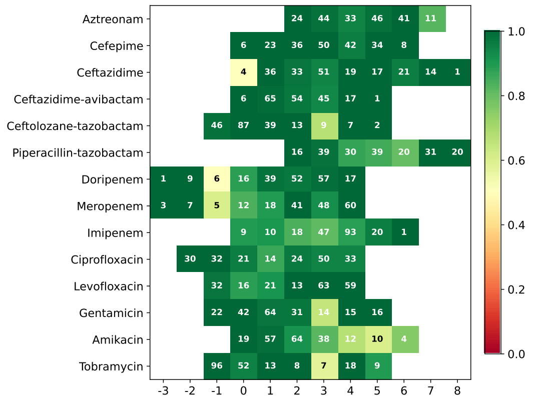


**Figure S2.** MICE all part precision miss 1

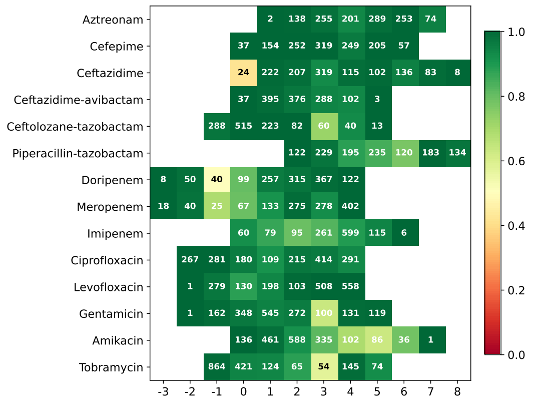


**Figure S3.** MICE all part precision miss 2

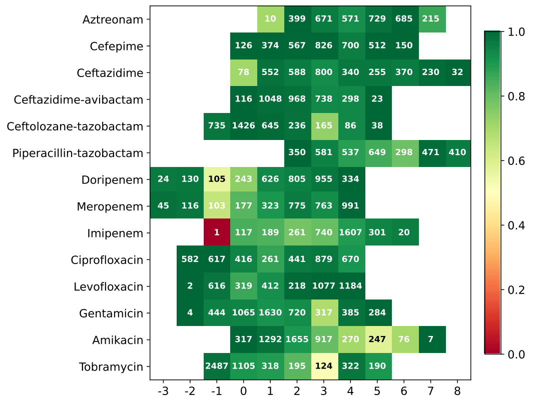


**Figure S4.** MICE all part precision miss 3

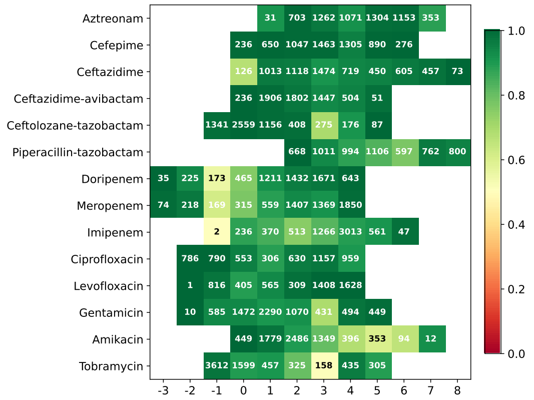

**Figure S5.** MICE all part precision miss 4

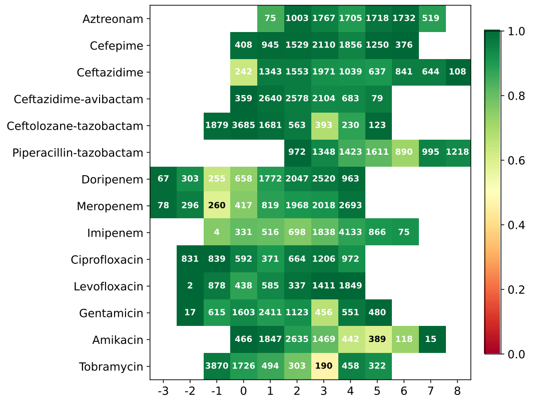

**Figure S6.** MICE all part precision miss 5

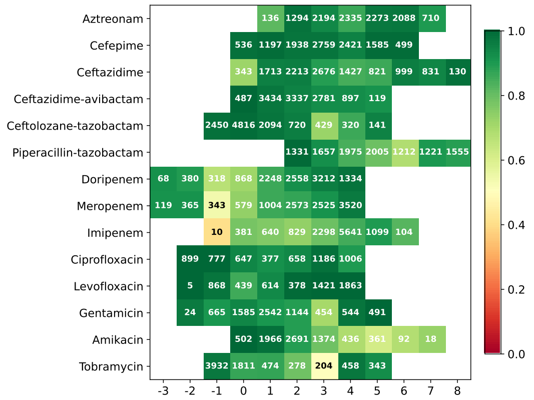

**Figure S7.** MICE all part precision miss 6

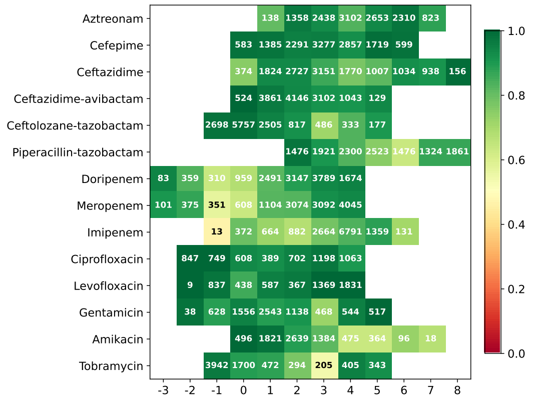

**Figure S8.** MICE all part precision miss 7

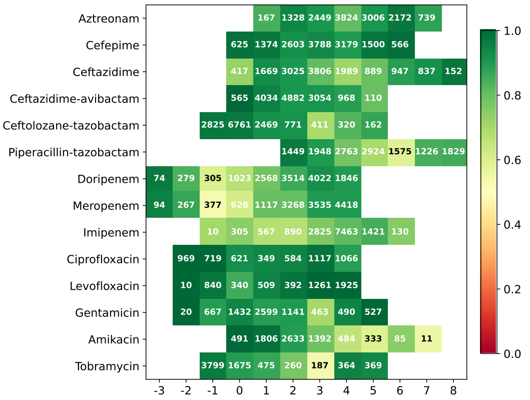


**Figure S9.** MICE all part precision miss 8

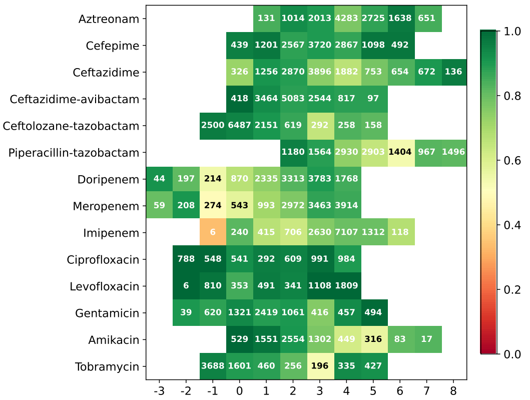


**Figure S10.** MICE all part precision miss 9

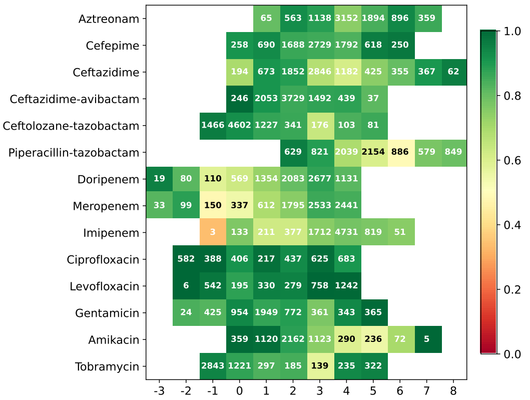


**Figure S11.** MICE all part precision miss 10

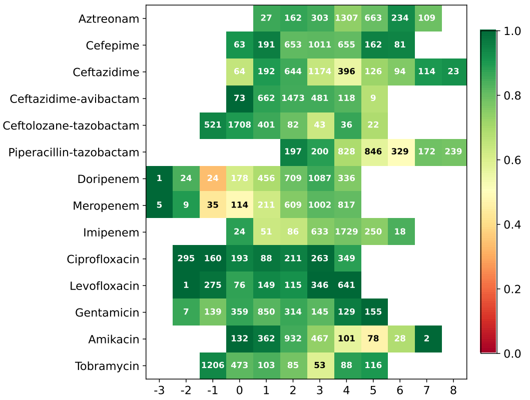


**Figure S12.** MICE all part precision miss 11

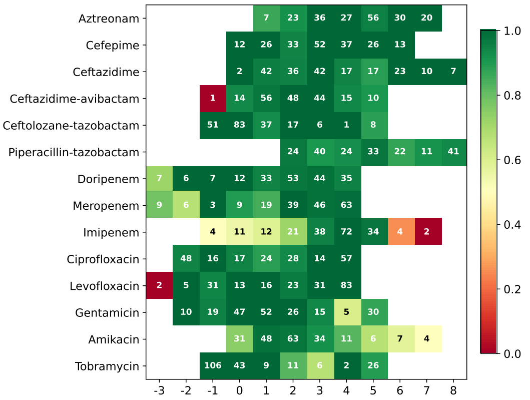


**Figure S13.** MICE all recall miss 1

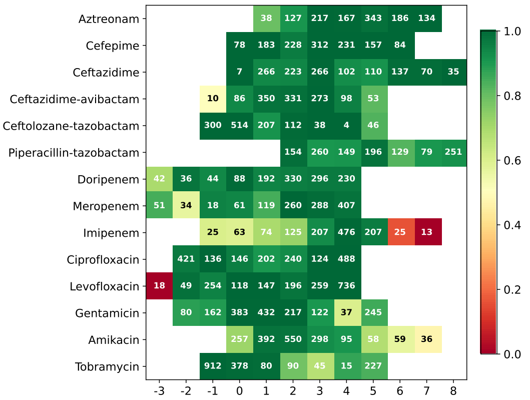


**Figure S14.** MICE all recall miss 2

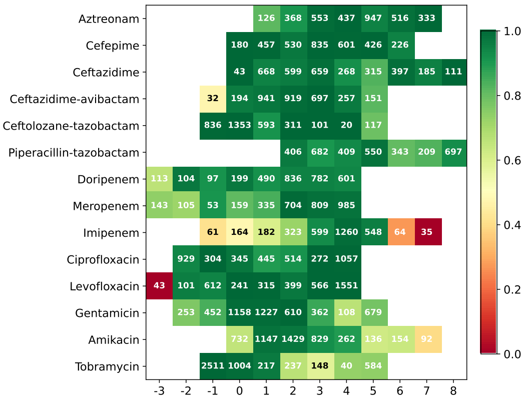


**Figure S15.** MICE all recall miss 3

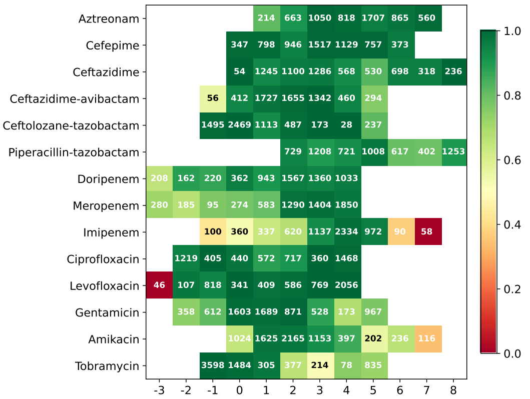


**Figure S16.** MICE all recall miss 4

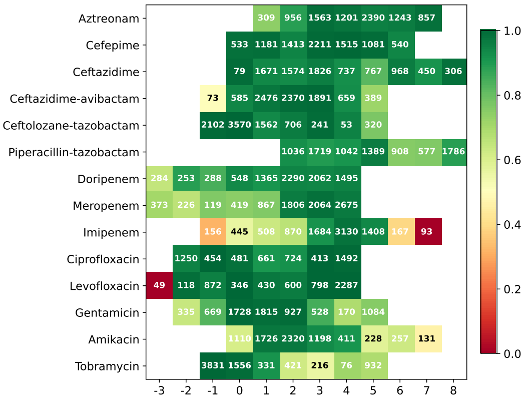


**Figure S17.** MICE all recall miss 5

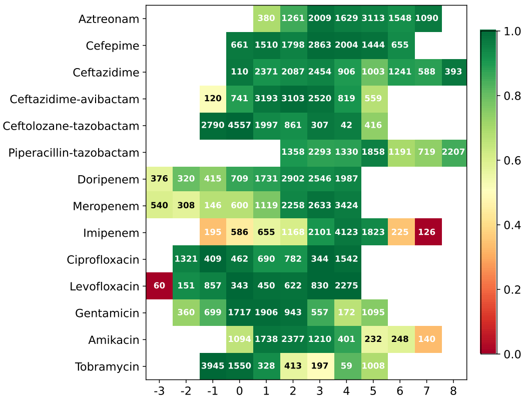


**Figure S18.** MICE all recall miss 6

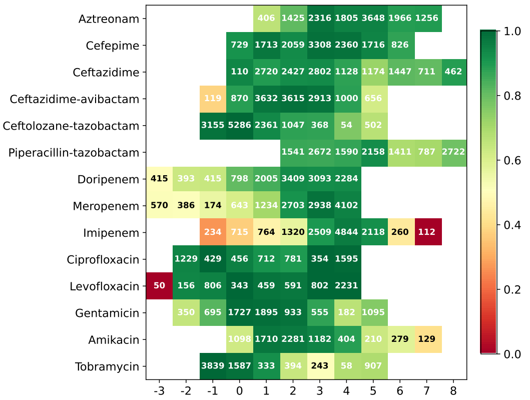


**Figure S19.** MICE all recall miss 7

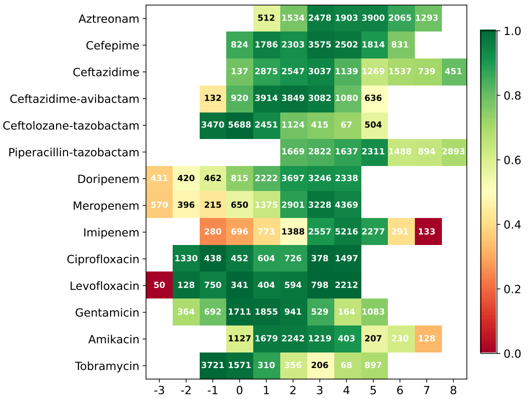


**Figure S20.** MICE all recall miss 8

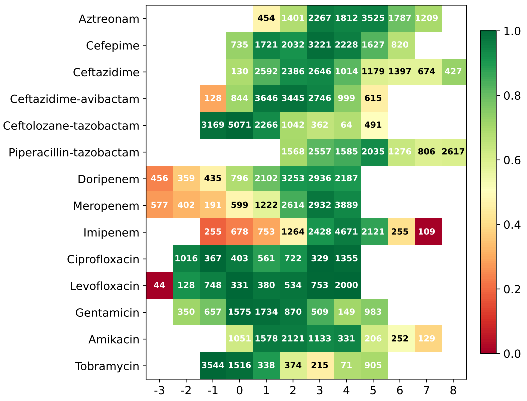


**Figure S21.** MICE all recall miss 9

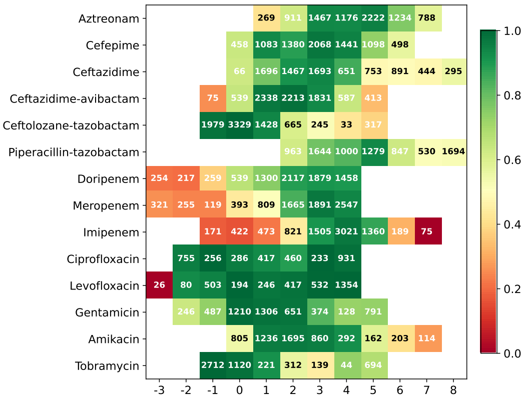


**Figure S22.** MICE all recall miss 10

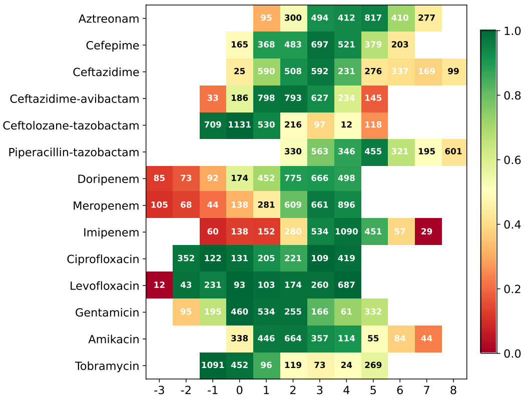


**Figure S23.** MICE all recall miss 11

### DAE performances

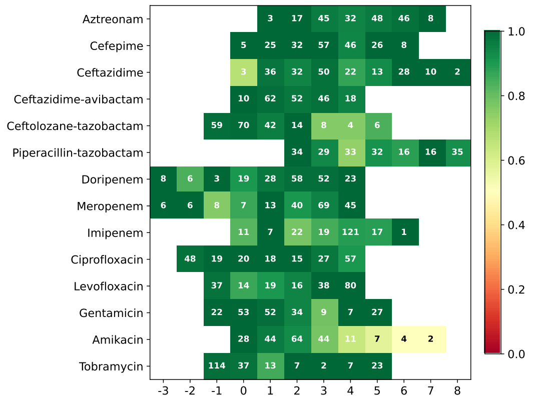


**Figure S24.** DAE all part precision miss 1

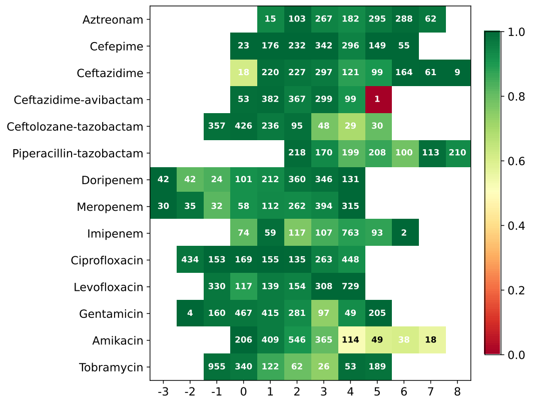


**Figure S25.** DAE all part precision miss 2

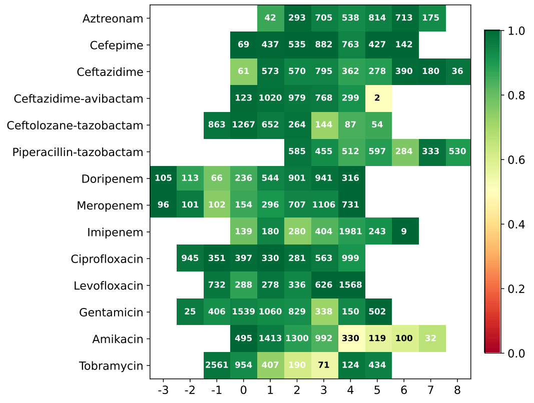


**Figure S26.** DAE all part precision miss 3

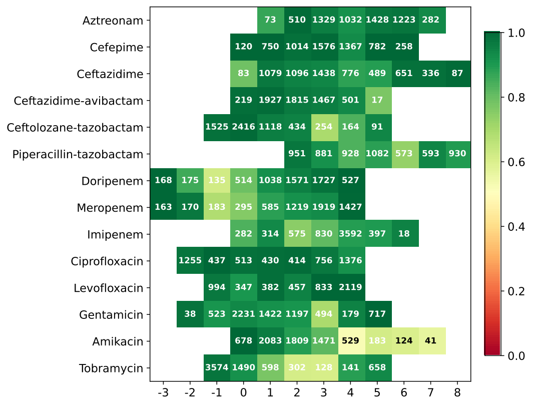

**Figure S27.** DAE all part precision miss 4

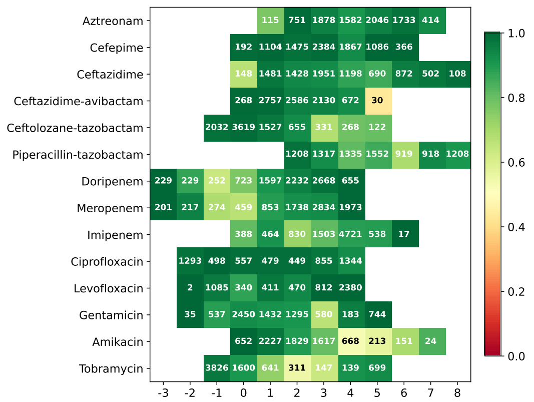

**Figure S28.** DAE all part precision miss 5

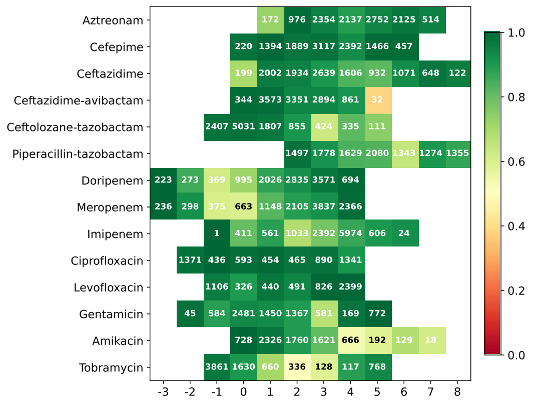

**Figure S29.** DAE all part precision miss 6

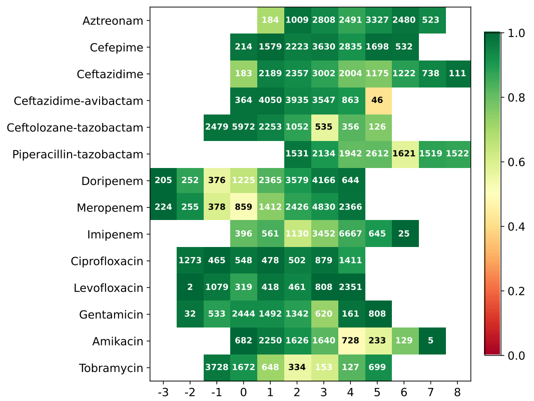

**Figure S30.** DAE all part precision miss 7

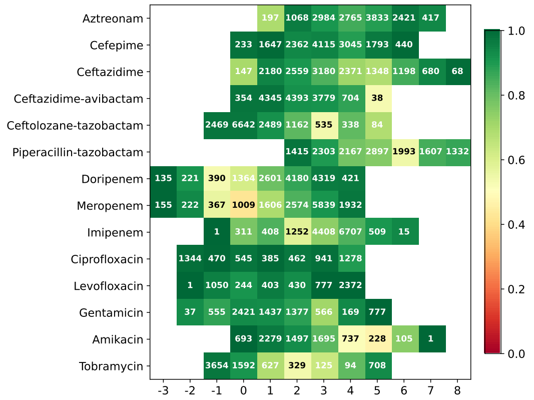

**Figure S31.** DAE all part precision miss 8

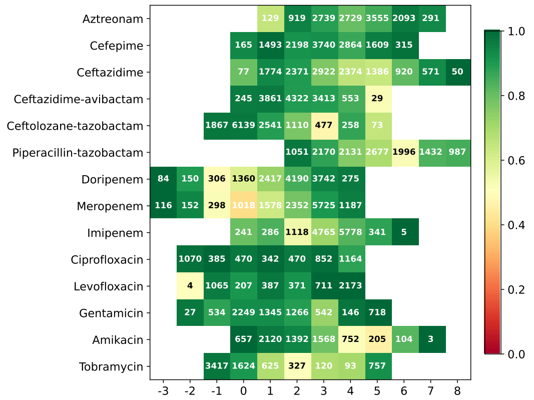

**Figure S32.** DAE all part precision miss 9

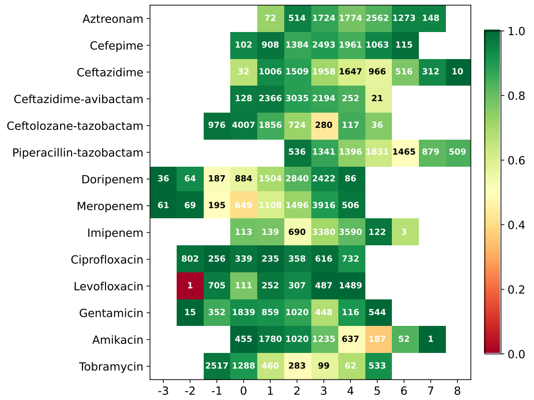

**Figure S33.** DAE all part precision miss 10

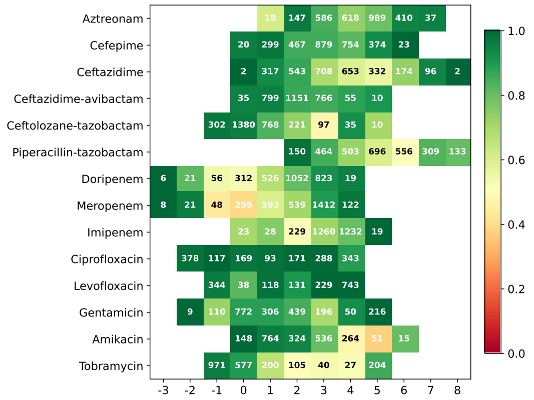

**Figure S34.** DAE all part precision miss 11

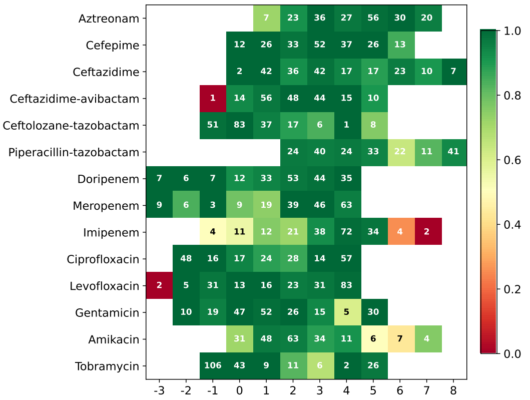


**Figure S35.** DAE all recall miss 1

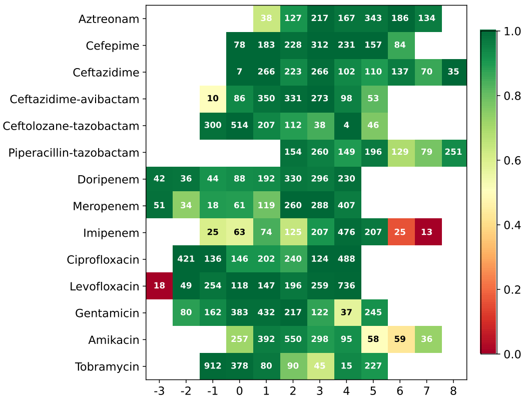


**Figure S36.** DAE all recall miss 2

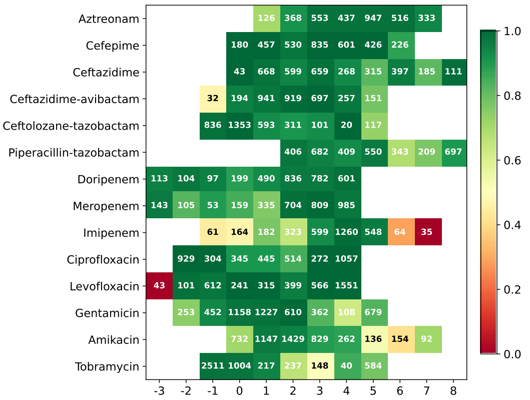


**Figure S37.** DAE all recall miss 3

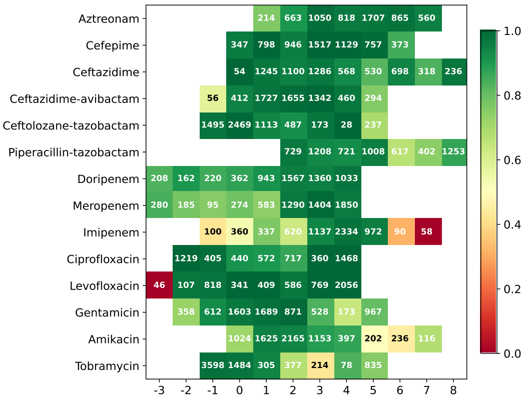


**Figure S38.** DAE all recall miss 4

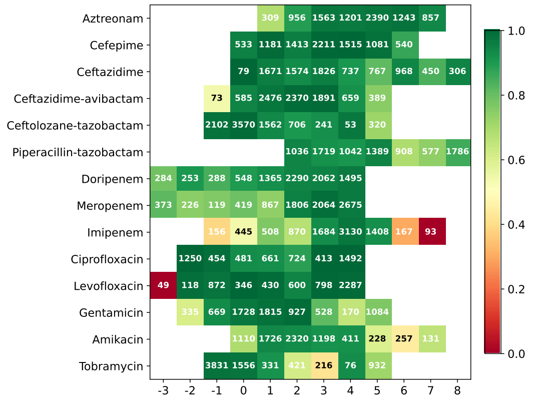

**Figure S39.** DAE all recall miss 5

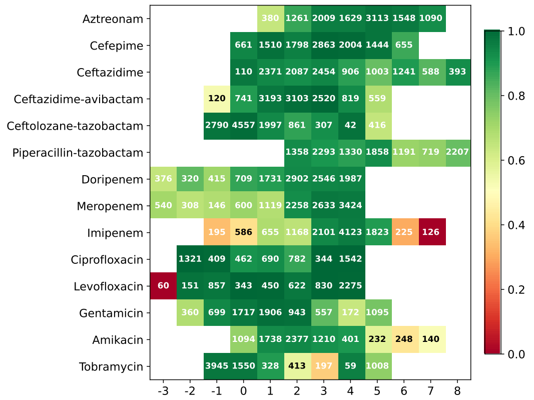

**Figure S40.** DAE all recall miss 6

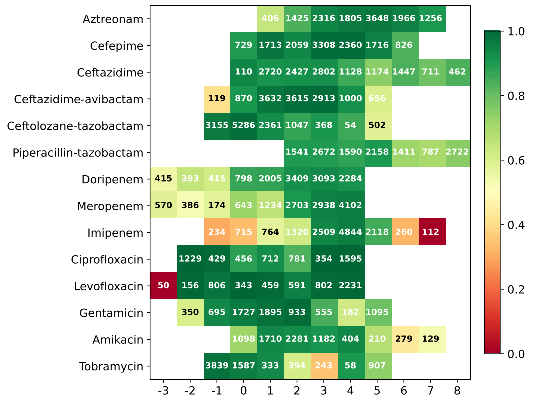

**Figure S41.** DAE all recall miss 7

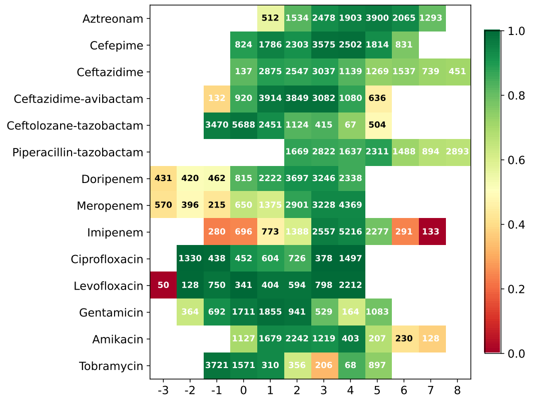

**Figure S42.** DAE all recall miss 8

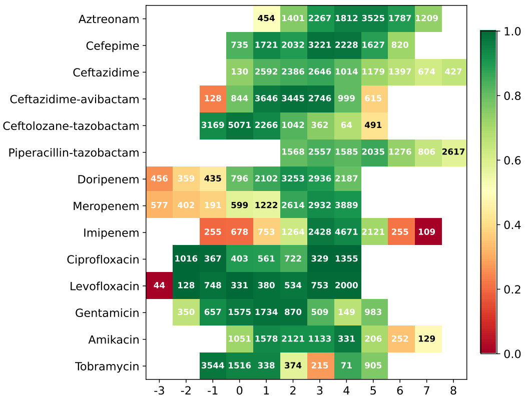


**Figure S43.** DAE all recall miss 9

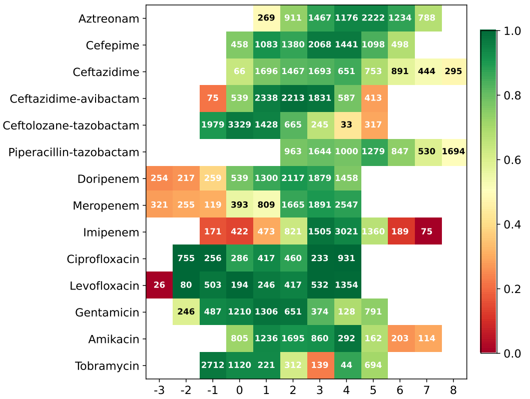


**Figure S44.** DAE all recall miss 10

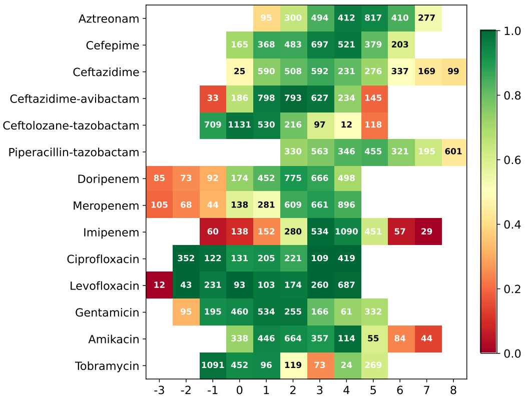


**Figure S45.** DAE all recall miss 11

#### References
